# Quantitative analysis of *in vivo* methionine oxidation of the human proteome

**DOI:** 10.1101/714089

**Authors:** John Q. Bettinger, Kevin A. Welle, Jennifer R. Hryhorenko, Sina Ghaemmaghami

**Affiliations:** Department of Biology, University of Rochester, NY, 14627, USA; University of Rochester Mass Spectrometry Resource Laboratory, NY, 14627, USA

**Keywords:** methionine oxidation, redox chemistry, hydrogen peroxide, oxidative protein damage, quantitative proteomics

## Abstract

The oxidation of methionine is an important posttranslational modification of proteins with numerous roles in physiology and pathology. However, the quantitative analysis of methionine oxidation on a proteome-wide scale has been hampered by technical limitations. Methionine is readily oxidized *in vitro* during sample preparation and analysis. In addition, there is a lack of enrichment protocols for peptides that contain an oxidized methionine residue; making the accurate quantification of methionine oxidation difficult to achieve on a global scale. Herein, we report a methodology to circumvent these issues by isotopically labeling unoxidized methionines with ^18^O labeled hydrogen peroxide and quantifying the relative ratios of ^18^O and ^16^O oxidized methionines. We validate our methodology using artificially oxidized proteomes made to mimic varying degrees of methionine oxidation. Using this method, we identify and quantify a number of novel sites of *in vivo* methionine oxidation in an unstressed human cell line.

## INTRODUCTION

The oxidation of methionine side chains has recently emerged as an important pathway for the posttranslational modification of proteins. Methionine is a normally hydrophobic amino acid with an oxidatively labile thioether group^1^. When oxidized, the chemical and physical properties of methionine are altered, causing alterations in protein structure and stability^2–4^. In this context, the oxidation of methionine has traditionally been thought of as a form of spontaneous protein damage. However, recent studies have shown that regulated oxidation of methionine can also serve as a mechanism for modulating protein function.

A notable example of regulatory methionine oxidation is carried out by a family of enzymes known as MICALs (molecule interacting with CasL), that enzymatically oxidize specific methionine residues on actin^5–7^. The oxidation of these methionine residues causes the formation of “fragile” F-actin filaments with an accelerated rate of depolymerization that has been shown to be important for the clearance of F-actin during cytokinesis and cell abscission^7–8^. Furthermore, it has been shown that the conformation-specific oxidation of methionine residues in calmodulin results in a reduced association between calmodulin and the plasma membrane and an increase in calmodulin degradation by the proteasome^9–10^. More recently, it has been shown that the liquid-liquid phase separation of the Ataxin-2 protein in *Saccharomyces cerevisiae* is regulated by the redox status of methionines in its low complexity domain^11^. In the case of F-actin, the oxidation of methionine is enzymatic, whereas in the latter two cases, it is believed to be chemical. Furthermore, methionine sulfoxides can be reduced by the action of specialized methionine sulfoxide reductase (Msr) enzymes, providing another potential mode of regulation^12–14^.

Despite its importance to protein structure and function, the large-scale investigation of methionine oxidation in a complex matrix, such as the cellular proteome, is hampered by technical limitations. Methionine oxidation has been shown to spuriously accumulate during the upstream stages of a typical bottom up proteomics workflow. In particular, methionine oxidation has been shown to increase with the length of trypsin digestion as well as the strength of ionization energy during electrospray ionization (ESI)^15–16^. These observations make it difficult to distinguish methionines that are oxidized *in vivo* from those that are artifactually oxidized *in vitro* during the course of sample preparation and mass spectrometric analysis. In addition, the bias of data dependent acquisition (DDA) for more abundant peptides and a lack of adequate enrichment protocols complicate the identification and quantification of methionines that are oxidized at low levels. Furthermore, oxidation of methionine residues results in significant changes in retention times and ionization propensities, making it difficult to accurately quantify fractional oxidation by comparing the relative intensities of oxidized and unoxidized spectra of methionine-containing peptides.

Several recent methods for the quantification of methionine oxidation have been developed with the aim of circumventing these technical limitations. Ghesquiere et. al demonstrated a method termed COFRADIC (combined fractional diagonal chromatography) proteomics^17^. This procedure isolates peptides that contain oxidized methionines by taking advantage of chromatographic shifts in reverse phase-high performance liquid chromatography (RP-HPLC) runs of peptides before and after the reduction of methionine sulfoxides by Msr enzymes. Using this methodology, the authors were able to isolate and identify a large set of oxidized methionine residues in a hydrogen peroxide stressed proteome from human jurkat cells. This method was successful in increasing the number of methionine sulfoxide containing peptides that were detected compared to traditional bottom up proteomic methods^18^. However, the COFRADIC approach requires multiple additional sample preparation steps prior to proteomic analysis as well as the production of an isotopically labeled refence proteome.

Liu et. al and Shipman et. al independently developed a strategy for the quantification of methionine oxidation that relies on the isotopic labeling of unoxidized methionine residues with H_2_^18^O_2_ during the early stages of sample preparation and prior to LC-MS/MS analysis^19–20^. This strategy results in the conversion of all unoxidized methionines to an ^18^O labeled version of the oxidized peptide. Conversely, peptides that are already oxidized *in vivo* retain their ^16^O modifications. The 2 Da mass difference between the ^16^O and ^18^O labeled methionine containing peptides is then used to distinguish between peptides that were unoxidized from those that were oxidized *in vivo*. The authors of these studies demonstrate that this strategy allows for the accurate quantification of methionine residues in a single protein.

Here, we report a modified version of the H_2_^18^O_2_ blocking methodology and extend the quantification of ^16^O/^18^O labeled methionine pairs to a proteome-wide level. Our strategy relies on the spectral identifications and MS1 annotations of the ^18^O labeled peptides, which are then used to identify, deconvolute and quantify the relative population of *in vivo* oxidized (^16^O-modified) peptides. We demonstrate the feasibility of this experimental approach and use it to measure *in vivo* methionine oxidation levels in unstressed human cells. Our data identifies a number of novel *in vivo* methionine oxidation sites while indicating that, as a whole, methionine oxidation is rare within the proteome of unstressed cells.

## EXPERIMENTAL PROCEDURES

### Cell culture, lysis and H_2_^18^O_2_ treatment

Wildtype human epidermal fibroblasts (MJT) cells were grown to confluency in Dulbecco’s Modified Eagle Medium (DMEM) supplemented with 15% Fetal Bovine Serum (FBS) and 1% penicillin-streptomycin (Invitrogen) and harvested by trypsinization. Cells were lysed in 50mM Triethylammonium bicarbonate (TEAB) (Fischer Scientific) and 5% sodium dodecyl sulfate (SDS) by high energy sonication and clarified of cell debris by centrifugation at 16,000xg for 10 minutes. Following lysis, protein concentration was quantified by Bicinchoninic Acid (BCA) assay and immediately diluted (1:1) to a final protein concentration of 0.5 mg/mL with either ^18^O heavy (Cambridge Isotope Laboratories) or ^16^O light (Fisher) H_2_O_2_ to a final H_2_O_2_ concentration of 1.25%. The oxidation reaction was allowed to continue for 1 hour at room temperature. Disulfide bonds were reduced by adding 2mM Dithiothreitol (DTT) (Fisher) and protein alkylation was performed with 10 mM iodoacetamide (IAA) (Sigma). Samples were acidified by adding phosphoric acid to a final concentration of 1.2% and subsequently diluted 7-fold with 90% methanol in 100mM TEAB. The samples were added to an S-trap column (Protofi) and the column was washed twice with 90% methanol in 100mM TEAB. Trypsin (Pierce) was added to the S-trap column at a ratio of 1:25 (trypsin:protein) and the digest reaction was allowed to continue overnight at 37°C. Peptides were eluted in 80 μL of 50 mM TEAB followed by 80 μL of 0.1% trifluoroacetic acid (TFA) (Pierce) in water and 80 μL of 50/50 acetonitrile/water in 0.1% TFA. Titration experiments were prepared by mixing light (^16^O) and heavy (^18^O) labeled proteomes in the specified ratio to a final protein amount of 50 μg. To increase proteome coverage, high-pH fractionation was conducted on extracts prior to LC-MS/MS. Extracts were fractionated using homemade C18 spin columns. Eight different elution buffers were made in 100 mM ammonium formate (pH 10) with 5%, 7.5%, 10%, 12.5%, 15%, 17.5%, 20%, and 50% acetonitrile added. All fractions were then lyophilized and re-suspended in 25 μl of 0.1% TFA.

### LC-MS/MS analysis

Fractionated peptides were injected onto a homemade 30 cm C18 column with 1.8 μm beads (Sepax), with an Easy nLC-1200 HPLC (Thermo Fisher), connected to a Fusion Lumos Tribrid mass spectrometer (Thermo Fisher). Solvent A was 0.1% formic acid in water, while solvent B was 0.1% formic acid in 80% acetonitrile. Ions were introduced to the mass spectrometer using a Nanospray Flex source operating at 2 kV. The gradient began at 3% B and held for 2 minutes, increased to 10% B over 5 minutes, increased to 38% B over 68 minutes, then ramped up to 90% B in 3 minutes and was held for 3 minutes, before returning to starting conditions in 2 minutes and re-equilibrating for 7 minutes, for a total run time of 90 minutes. The Fusion Lumos was operated in data-dependent mode, with MS1 scans acquired in the Orbitrap, and MS2 scans acquired in the ion trap. The cycle time was set to 1.5 seconds to ensure there were enough scans across the peak. Monoisotopic Precursor Selection (MIPS) was set to Peptide. The full scan was collected over a range of 375-1400 m/z, with a resolution of 120,000 at m/z of 200, an AGC target of 4e5, and a maximum injection time of 50 ms. Peptides with charge states between 2-5 were selected for fragmentation. Precursor ions were fragmented by collision-induced dissociation (CID) using a collision energy of 30% with an isolation width of 1.1 m/z. The ion trap scan rate was set to Rapid, with a maximum injection time of 35 ms, and an AGC target of 1e4. Dynamic exclusion was set to 20 seconds

### MaxQuant analysis and data conversion

Raw files for all samples were searched against the H. sapiens Uniprot database (downloaded 8/7/2017) using the integrated Andromeda search engine with MaxQuant software. Peptide and protein quantification were performed with MaxQuant using the default parameter settings. ^18^O methionine Sulfoxide, ^16^O methionine Sulfoxide, ^18^O methionine Sulfone and N-terminal acetylation were set as variable modifications and carbamidomethyl cysteine was set as a fixed modification. Raw files were converted to mzXML format with the msconverter software using the vendor supplied peak picking algorithm and the threshold peak filter set to the top 1500 peaks in each scan. The MaxQuant supplied evidence file and the mzXML file were used as input into a custom algorithm described below.

When analyzing the prevalence of other H_2_O_2_-induced modifications (Figure 2B), Maxquant searches were conducted for unmodified peptides (except for carbamidomethylation of cysteines as a constant modification). The total measured intensities of all unmodified peptide spectral matches (PSMs) containing specific residues were measured as fractions of total intensities of all PSMs.

### Custom search and quantification

For each MS1 peptide feature that was identified by MaxQuant, a theoretical isotope cluster was generated using the atomic composition of the peptide and the known abundance of natural isotopes. Next, for each peptide feature, the associated retention time range identified by MaxQuant was used to subset a swath of the MS1 spectra predicted to contain the associated feature. The swath was then filtered for peaks with a centroided mass to charge ratio within a 14 ppm window centered on the predicted mass to charge ratios for the identified peptide feature. Assembled peaks were then summed across the retention time range and projected onto two separate 2D planes: the m/z-intensity plane and the retention time-intensity plane. Isotopic peaks were sequentially connected along the m/z axis only if they improved the fit to the theoretical isotope cluster along the m/z-intensity plane.

The spectra were then mined for the differentially labeled peptide within the same retention time window. If the identified peptide was unlabeled, the search defaulted to searching for the light (−2Da) modification within the retention time window. Using the atomic composition of the peptide theoretical models were generated for each peptide with theoretical ratios between the light and heavy labeled peptide ranging between 0 and 1 with a step size of 0.01. The theoretical model with the best fit to the observed data along the m/z-intensity plane was taken as the measured light:heavy ratio for that peptide.

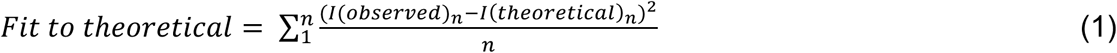

Where I_n_ is the summed intensity of each isotopic peak and n is the number of isotopic peaks assembled into the model. Intensities were expressed as values ranging between 0 and 1 normalized by the most intense isotopic peak.

Three measures of model quality were used to asses each model and assign statistical confidence. On the retention time-intensity plane, a spearman correlation in the retention time profiles between the light and heavy labeled peptide was measured (referred to as RT Spearman Correlation in Figure 3). In addition, a percent shift in the center of masses of the elution profiles between the heavy and light labeled peptide was measured (referred to as RT Shift in Figure 3).

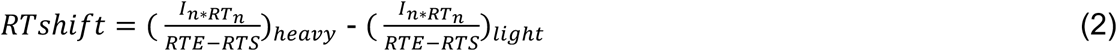

Where I_n_ is the intensity of a datapoint expressed as a value ranging between 0 and 1 normalized by the maximum intensity and RT_n_ is the associated retention time. Next, the accumulated error along the m/z axis of the individual data points used to create the model was measured as the average difference between the theoretical mass to charge ratio and the centroided mass to charge ratio of the peaks assembled into the model (referred to as m/z error in Figure 3).

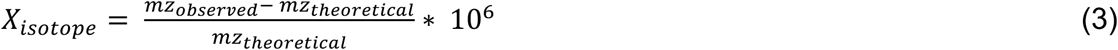

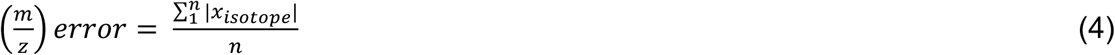

Where n is equal to the number of isotopic peaks assembled into the model. Within each isotopic peak the negative and positive m/z errors were allowed to cancel each other in order to maximize sensitivity to mass drift. However, between isotope clusters, the absolute value of the error is averaged.

Each peptide feature has a uniquely quantified model, however most peptides are represented by multiple peptide features. As a final step, peptide features were aggregated together according to their sequence and modification status. Measurements taken along the m/z axis (fraction oxidized and m/z error) were reported as the intensity weighted mean of each feature associated with the peptide.

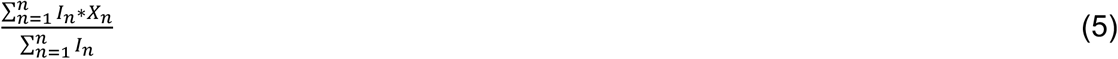

Where I_n_ is equal to the total intensity of the assembled model and X_n_ is equal to the fraction oxidized or m/z error associated with the model. Measurements taken along the retention time axis (Spearman correlation, relative RT shift) were reported as the cumulative result after consolidating all features associated with a peptide into a single feature. As a final step, the measured fraction oxidize for each peptide was corrected by the labeling efficiency of that peptide. Labeling efficiencies were calculated by comparing the assembled intensity of the oxidized peptide to the unoxidized version.

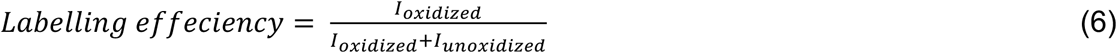

All subsequent data analysis was restricted to peptides with a 90% labeling efficiency or greater.

### Assigning Statistical Confidence

Unlabeled peptides are not expected to have a −2Da modification and the model assembled for those peptides are an experimental approximation of the null distribution that is measured as a function of background alone. All (-) methionine, (-) cysteine containing peptides are unlabeled and assigned to a decoy dataset, whereas all (+) methionine, (-) cysteine are assigned to a target dataset Quality parameters discussed above are reduced to a single score by principal component analysis. The distribution of principle component scores within the decoy dataset (unlabeled peptides) is then used to calculate a false positive rate (FPR) for each score.

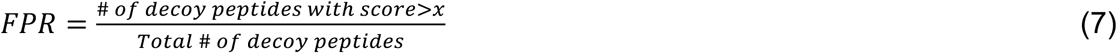

## RESULTS

### ^18^O blocking methodology

We developed a stable isotope labeling strategy that allows for the accurate quantification of *in vivo* methionine oxidation while preventing artifactual *in vitro* methionine oxidation that typically occurs during the course of a bottom-up proteomic workflow. An overview of the experimental strategy is illustrated in Figure 1. The spontaneous *in vitro* oxidation of methionine during sample preparation may result in an overestimation of *in vivo* oxidation levels. We circumvent this problem by forced oxidation of methionines with ^18^O labeled hydrogen peroxide (H_2_^18^O_2_) at the time of cell lysis. Hence, relative intensities of ^16^O and ^18^O-modified methionine-containing peptides provides a measure of *in vivo* fractional oxidation prior to lysis.

**Figure 1.**
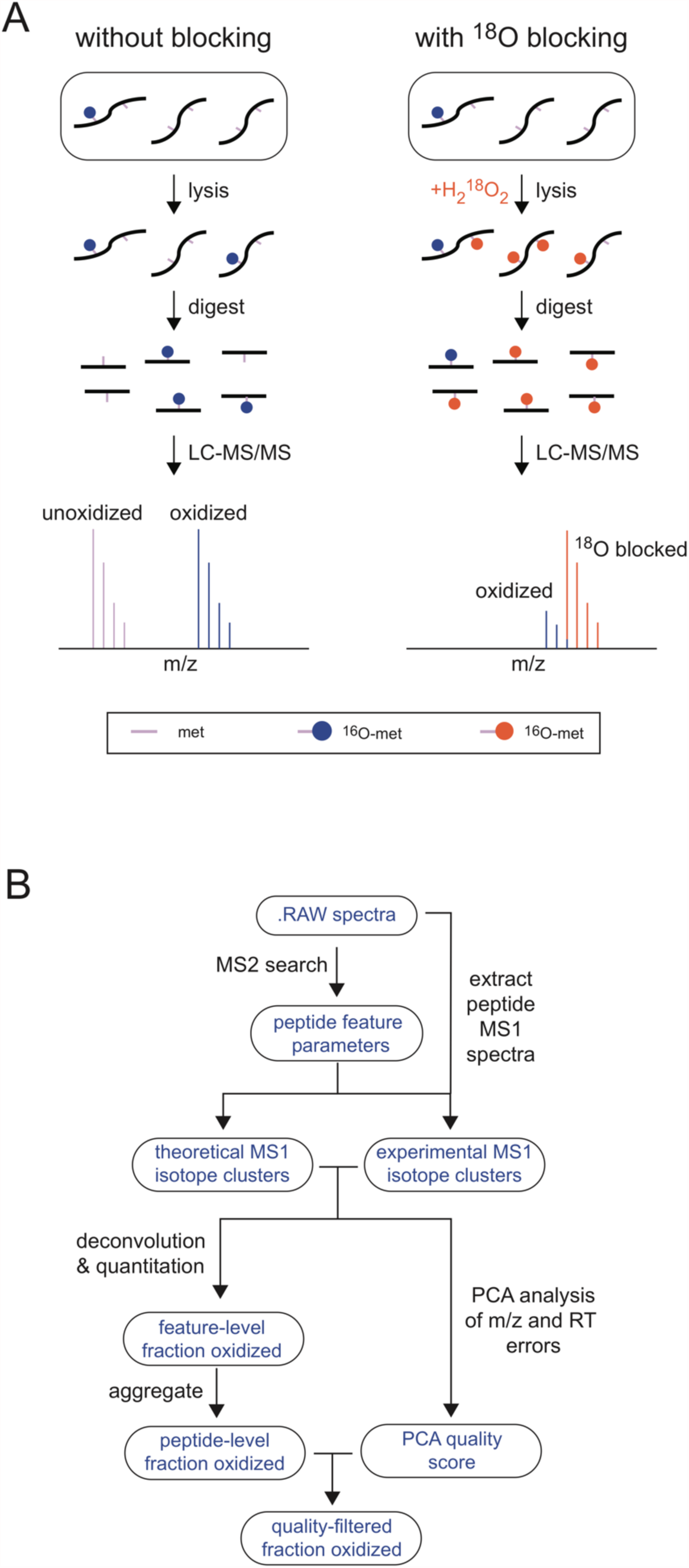
Schematic overview of ^18^O blocking methodology. (A) A comparison of strategies for quantifying methionine oxidation with (left) or without (right) blocking unoxidized methionines with a heavy labeled oxidizing agent. Unlike blocked samples, unblocked samples accumulate oxidized methionines during sample preparation, resulting in an overestimation of oxidation levels. (B) A general overview of the computational strategy for finding, deconvoluting and quantifying the relative ratios of light and heavy labeled peptides in a blocked samples (see Experimental Procedures for additional details).

To rapidly block *in vivo* unoxidized methionines with ^18^O, cells were lysed in a denaturing buffer and subsequently exposed to excess levels of ^18^O labeled hydrogen peroxide. Oxidation was allowed to proceed for 1 hour at room temperature. These oxidation conditions allows for the near complete labeling of all methionine containing peptides (Figure 2A). In oxidized extracts, only ∼5% of the total methionine containing peptide intensities were due to unlabeled (unoxidized) peptides, <1% were due to doubly oxidized methionine sulfones, and the remaining ∼95% were due to light or heavy labeled methionine sufloxides (Figure 2A). Independently conducted searches showed a low prevalence of other oxidized residues with the exception of cysteines (Figure 2B). Because of this potential complication, peptides containing both methionines and cysteines (∼1% of all methionine-containing peptides) were excluded from analysis.

**Figure 2.**
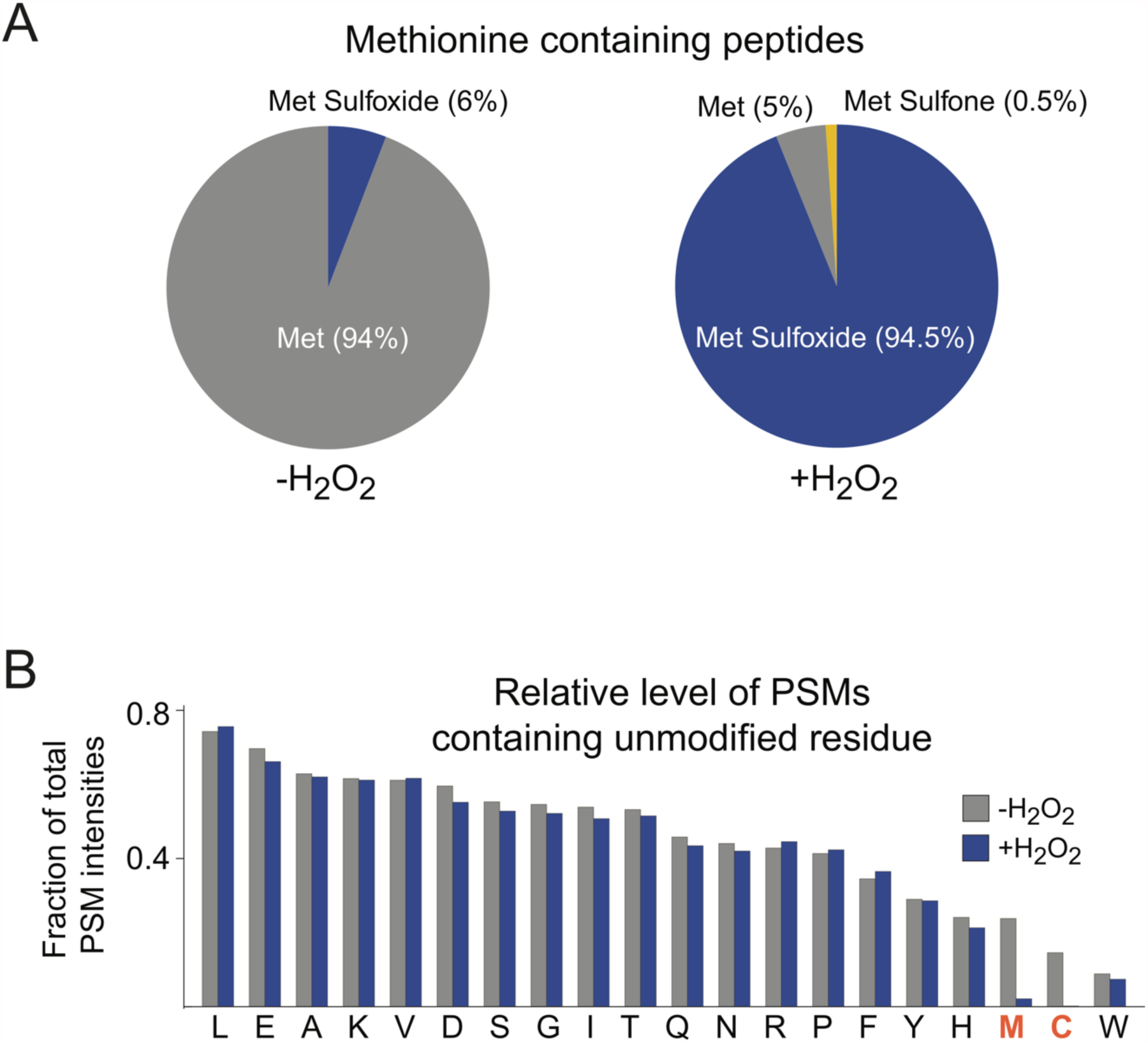
Validation of the oxidation conditions. (A) The total spectral intensity of single methionine containing peptides with no cysteines in unblocked samples (left) are predominately composed of peptides with unoxidized methionines with a small fraction of peptides containing methionine sulfoxides. This trend is reversed in blocked samples (right) where the predominant variants are methionine sulfoxide containing peptides with a small fraction of unoxidzed peptides and peptides with doubly oxidized methionine sulfones. (B) Comparison of relative numbers of PSMs found for unmodified peptides containing at least one of each amino acid both before and after blocking with hydrogen peroxide. Amino acids that are found to be significantly modified by hydrogen peroxide treatment are shown in red.

### Quantitation of fractional oxidation

A full description of the data analysis pipeline is provided in Experimental Procedures and a brief graphical overview is shown in Figure 1B. Since *in vivo* ^16^O-oxidized methionines were rare within the unstressed MJT cells (see below), ^18^O modified forms of methionine-containing peptides were typically identified by mass spectrometric data dependent acquisition (DDA) of ^18^O-oxidized samples. The measured retention time (RT) and mass to charge ratios (m/z) of the ^18^O labeled peptides were then used to conduct a model dependent search to identify the entire isotope cluster and measure the relative ratios of ^16^O and ^18^O-modified variants of the peptide. Since ^16^O and ^18^O-modified peptides differ in mass by only 2 Da, quantitation of their relative levels requires deconvolution of isotopic clusters. In rare cases where the ^16^O labeled version of methionine-containing peptides were identified by DDA, the quantitation was reversed to maximize coverage.

MS1 spectra of crude extracts are generally complex and light ^16^O labeled peptides commonly have low abundance relative to background peptides. We were therefore concerned that some of our assembled isotope cluster models may be dominated by background signal coming from overlapping peptide features. In order to distinguish models representing true ^16^O labeled peptides from background, we calculated a quality score for each model that was dependent on the chromatographic correlation between the ^16^O and ^18^O labeled peptides as well as the m/z error of individual peaks assembled into the model. To assess the efficacy of our scoring function to distinguish true hits from background, we conducted a decoy search composed of unlabeled peptide spectra for which we did not expect a true −2 Da modification (e.g. non-methionine containing peptides). Figure 3 shows a comparison of the distribution of scores calculated for labeled vs unlabeled peptides in our baseline titration point. The calculated scores show strong correlations with the different measures of model error (Figure 3B) and labeled peptides have a significantly higher distribution of scores than unlabeled peptides (Figure 3A). This suggests that our scoring strategy can be used to distinguish true hits from background. Using the distribution of scores in the unlabeled dataset, a false positive rate (FPR) was calculated for each score. All subsequent data analysis is restricted to peptides with scores associated with an FPR of <10%. An additional filter is applied to the fit of the theoretical isotope cluster with a hard cutoff of 0.005. The filter applied to the fit to the theoretical isotope cluster is a user set parameter and was adjusted in order to balance accuracy and sensitivity. Together, the data in Figure 3 indicate that our scoring strategy effectively limits the level of background noise in our dataset and allows us to identify highly confident models.

**Figure 3.**
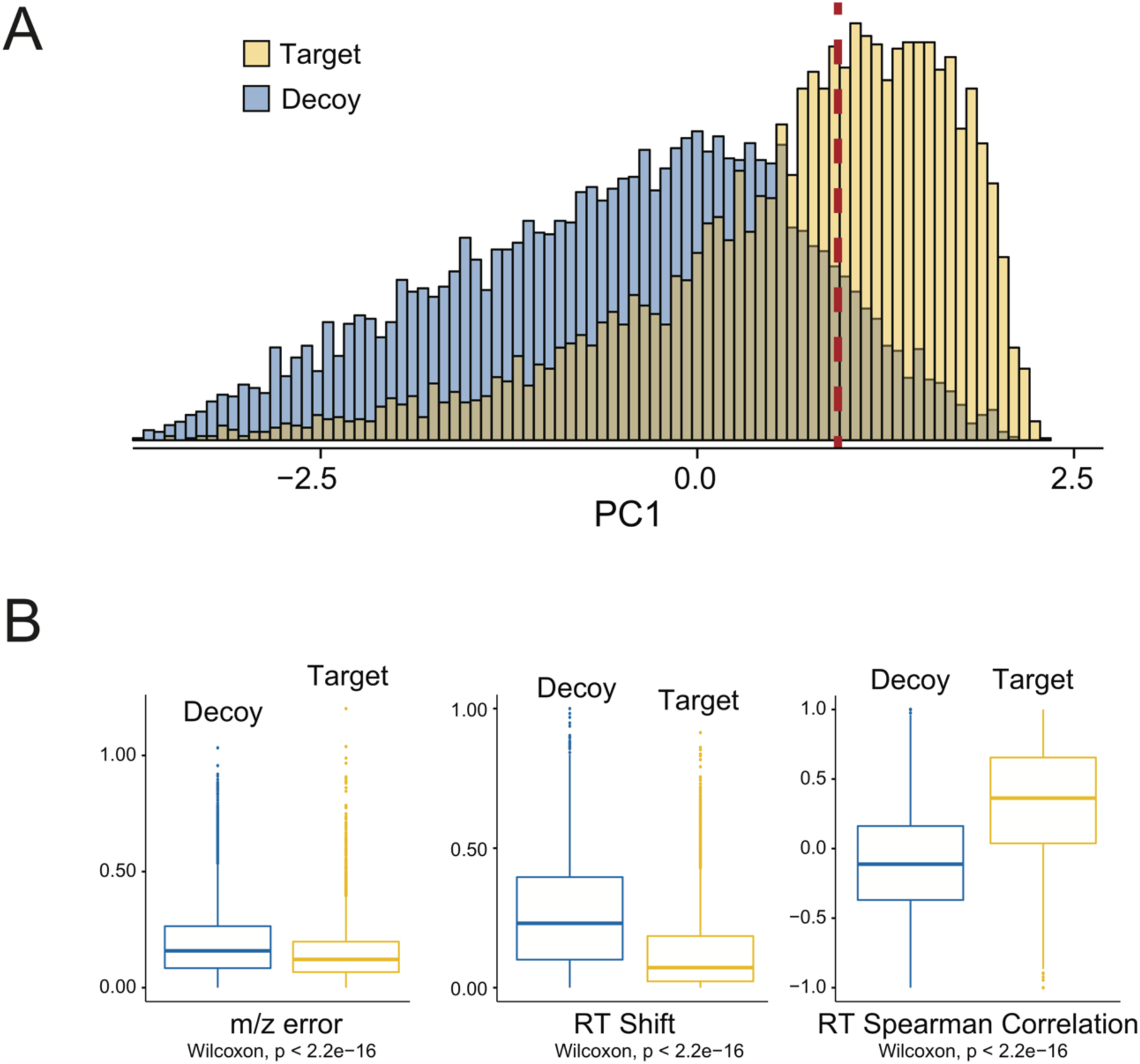
Quality control scoring results. (A) Principle component analysis (PCA) of the quality parameters shown in (B) was used to calculate a quality score for each peptide. Methionine-containing peptides (Target) for which a true −2 Da modification is expected to exist have better scores than non-methionine-containing peptides peptides (Decoy) for which no −2 Da modification is expected to exist. The red dotted line shows the principle component value at which there is a global 10% false positive rate (FPR). All subsequent data analysis is done on peptides with principle component scores that are better than the cutoff value represented by the red dotted line. (B) The three quality parameters used to calculate the principle component score (see Materials and Methods) show significant distinction between Target and Decoy peptides. For both (A) and (B), results from the baseline titration point (0.0 ^18^O : 1.0 ^16^O) is shown as a representative example.

### Quantitative validation of fractional oxidation measurements

As a first application, we sought to validate our quantitative approach by conducting a spike-in titration experiment. At varying ratios, fully light (^16^O) labeled proteome prepared from MJT cells was spiked into a background of a heavy (^18^O) labeled proteome from the identical cell extract. Four titration samples were prepared with spike-in ratios of 0:1.0, 0.1:0.9, 0.25:0.75 and 0.4:0.6. As expected, the measured distribution of light:heavy ratios increase proportionally in accordance to spike-in ratios (Figure 4, Supplementary Table 1). Importantly, the measured light:heavy ratios for the decoy dataset does not increase with the amount of labeled spike-in, suggesting that they represent a true measurement of background (Figure 4B, Supplementary Table 1).

**Figure 4.**
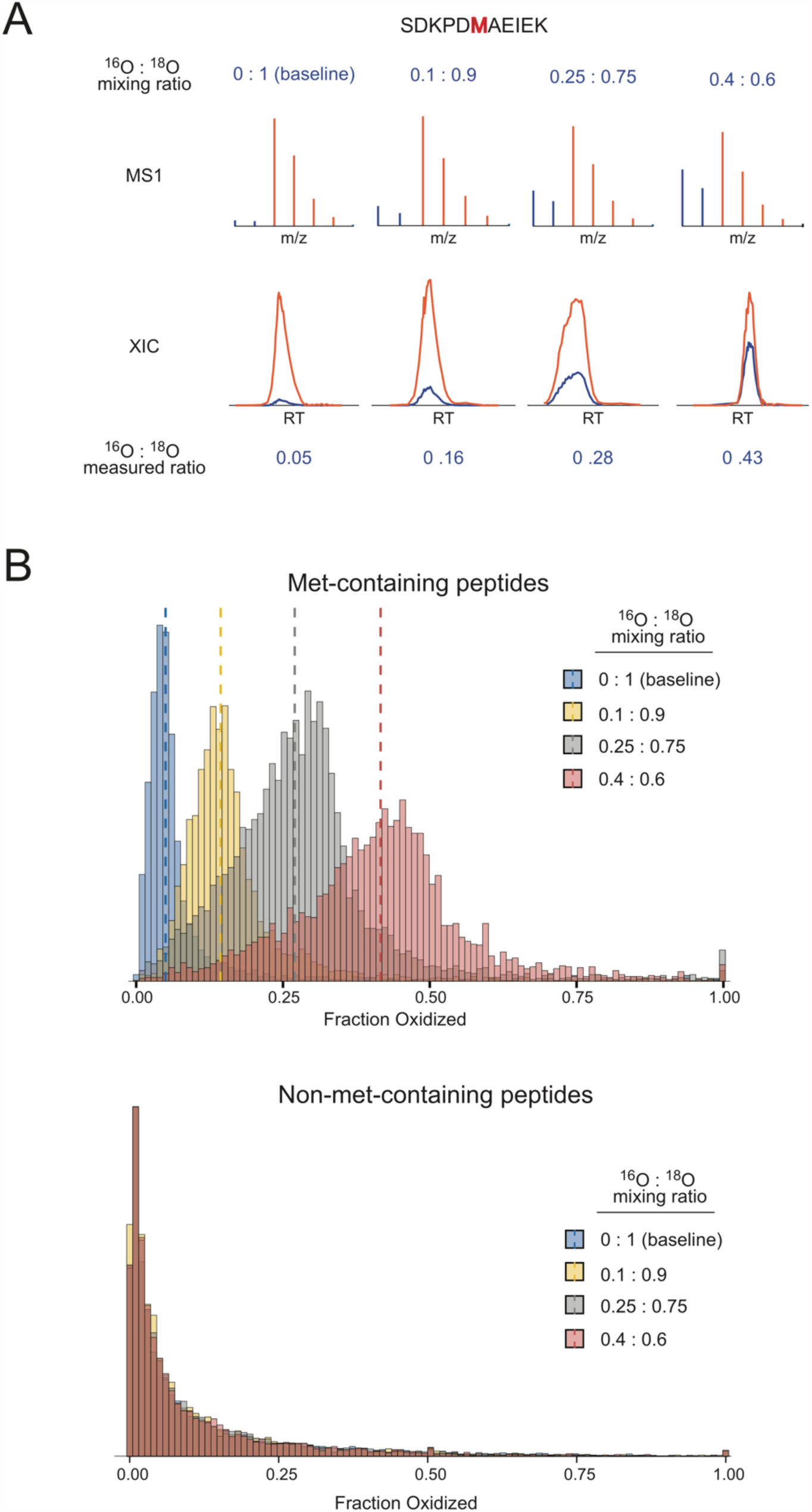
Spike-in titration results and validation of methodology. (A) A peptide detected in all titration points is shown as a representative example. The MS1 spectra of one feature representing the peptide is shown at the top and the total XIC of all features associated with the peptide is shown at the bottom. Red and blue colors represent the peaks and intensities corresponding to ^18^O and ^16^O peptides, respectively. The top and bottom row of numbers indicate the mixing ratio and the measured fractional oxidation, respectively. (B) The distribution of measured fractional oxidation levels of all single methionine-containing peptides with no cysteines that passed the quality score filter is show on top. Dotted lines indicate the respective median values. The null distribution of fractional oxidation levels for all non-methionine containing peptides is shown at the bottom.

As tabulated in Table 1, the measured medians of fractional oxidation for the four spike-in experiments were 0.050, 0.140, 0.280 and 0.421, respectively. It is important to note that although the spike-in sample can be assumed to be fully light (^16^O) labeled, the same cannot be assumed of the heavy labeled sample. The heavy labeled sample was not cleared of all *in vivo* (^16^O) methionine oxidation prior to labeling, and the H_2_^18^O_2_ used as the labeling reagent contains some ^16^O isotopic impurity, thus the heavy labeled background sample contains a mixture of ^16^O and ^18^O labeled methionine containing peptides. We therefore calculated the expected median for each titration point using a dilution formula that takes into account the measured median of the baseline titration point as a value representative of the amount of light labeled peptides present in the background prior to spike-in. For example, given that the measured median of the baseline titration point was 0.051, in the 0.1:0.9 titration point we would expect a measured fractional oxidation of 0.14 (0.1×1 + 0.9×0.05). Table 1 lists the measured medians, and the expected median. The difference between the expected median and the measured median did not exceed a maximum value of 0.9%, indicating that our experimental approach is capable of measuring the global levels of methionine fractional oxidation levels with a high degree of accuracy.

**Table 1.**
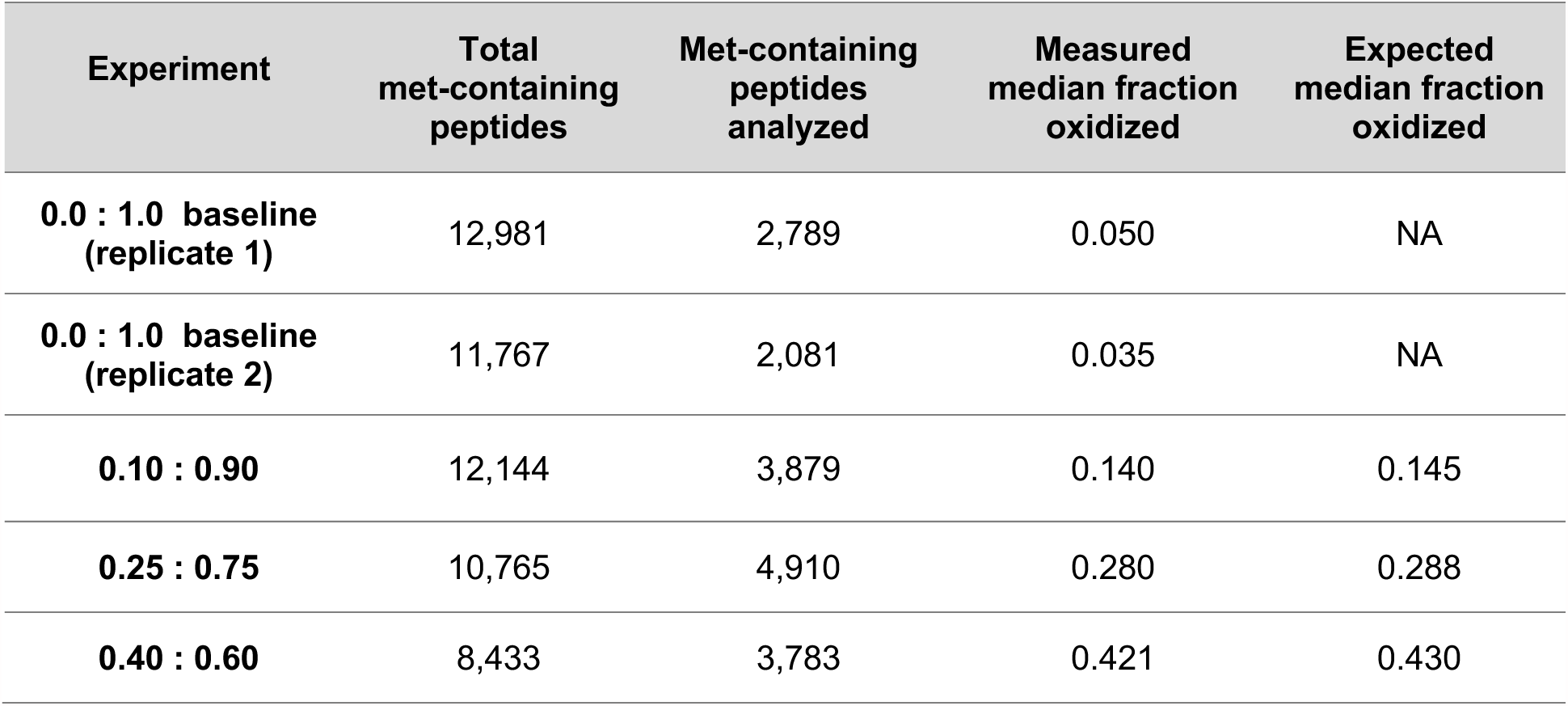
Global statistics of methionine oxidation

### Levels of *in vivo* methionine oxidation

The distribution of methionine oxidation levels in an unstressed human proteome appears to be significantly lower than previous estimates made on oxidatively stressed human proteomes^17^. Within our baseline measurement of methionine oxidation in an unstressed human proteome, 39.8% of methionine containing peptides had levels of *in vivo* oxidized (light labeled) peptides that were below the limit of detection. The levels of light labeled peptides that were below the limit of detection decreased as the amount of light labeled spike increased (Table 1). For those peptides that had detectable levels of methionine oxidation, the median value measured for fractional oxidation was 5.0%, a value that is within the limit of isotopic impurity of H_2_^18^O_2_ provided by the manufacturer (Figure 5).

**Figure 5.**
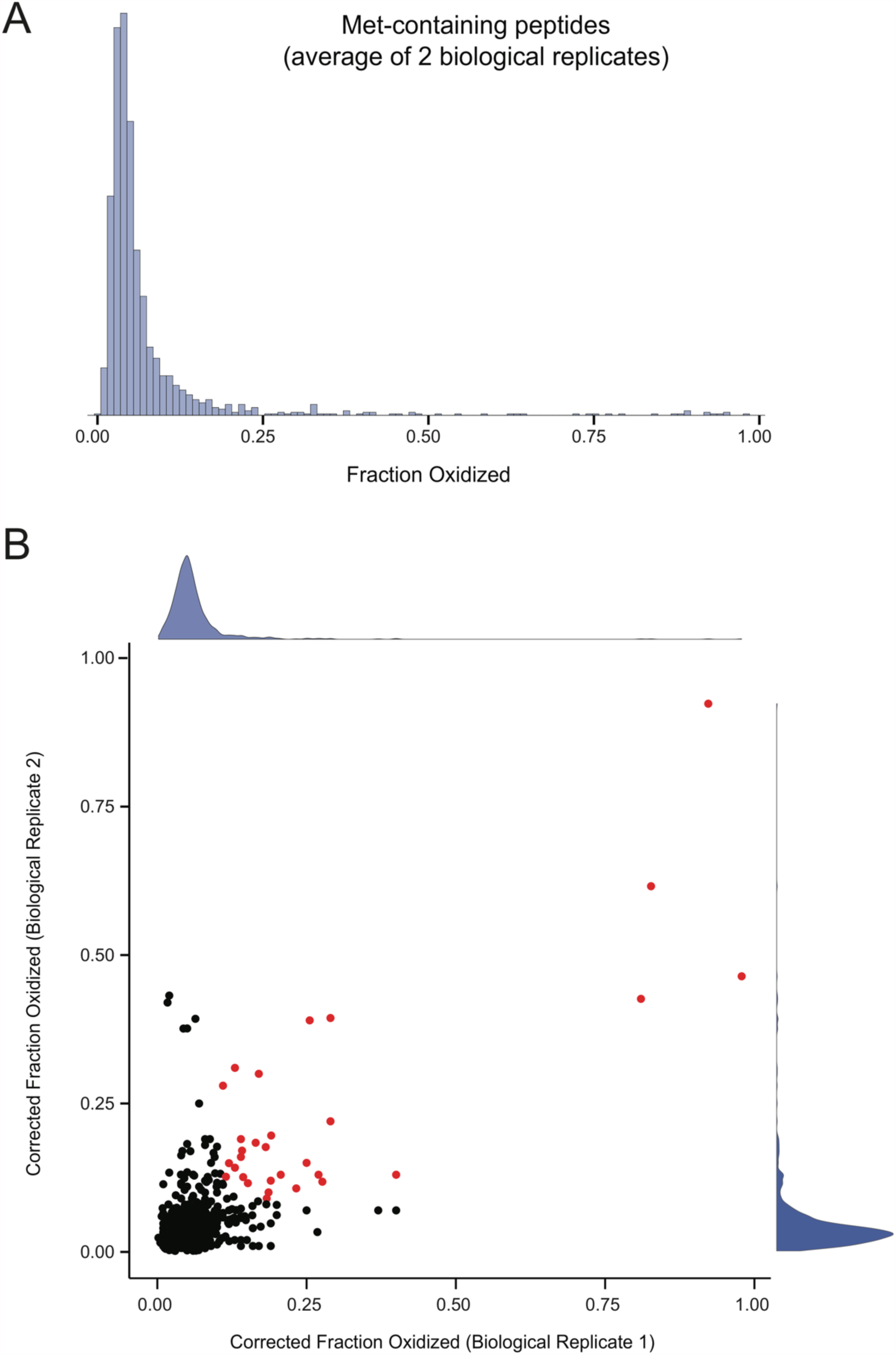
The distribution of *in vivo* methionine oxidation levels in unstressed MJT cells. (A) Measurements of methionine oxidation levels were made in two biological replicates and the distribution of the averaged results are shown. (B) A comparison of peptide level measurements of methionine oxidation between two biological replicates. The distributions of each replicate are shown on the respective axes. Dots are colored red if they are found to be highly and reproducibly oxidized in both replicates and have a labeling efficiency of at least 80% in both replicates. Peptides represented by the red colored datapoints are tabulated in Supplementary Table 2.

Measurements conducted on a biological replicate confirmed the observation that the levels of methionine oxidation in an unstressed human proteome are low and within the limit of isotopic impurity with a median value of 3.5% (Figure 5, Table 1). We observed a a slight shift in observed methionine oxidation levels between the biological replicates. However, biological replicates were prepared using separate batches of ^18^O labeled hydrogen peroxide, making it difficult to ascribe the observed shift to a biological source and highlighting the importance of batch effects in our assay. Nonetheless, we observed good agreement in the measured levels of methionine oxidation between replicates on a peptide to peptide level and were able to qualitatively detect a subset of peptides that were significantly oxidized in both replicates. (Figure 5B).

For a subset of the isotope clusters assembled and used to quantify methionine oxidation, there is direct overlapping MS/MS evidence that both labeling variants of the peptide are present in the model (Supplementary Table 2). Despite the observation that global levels of methionine oxidation are generally low, we observed a subpopulation of methionine containing peptides that are observed to have reproducibly high levels of methionine oxidation *in vivo* (Supplementary Table 2). A representative example of a verified and highly oxidized peptide is shown in Figure 6 and discussed below.

**Figure 6.**
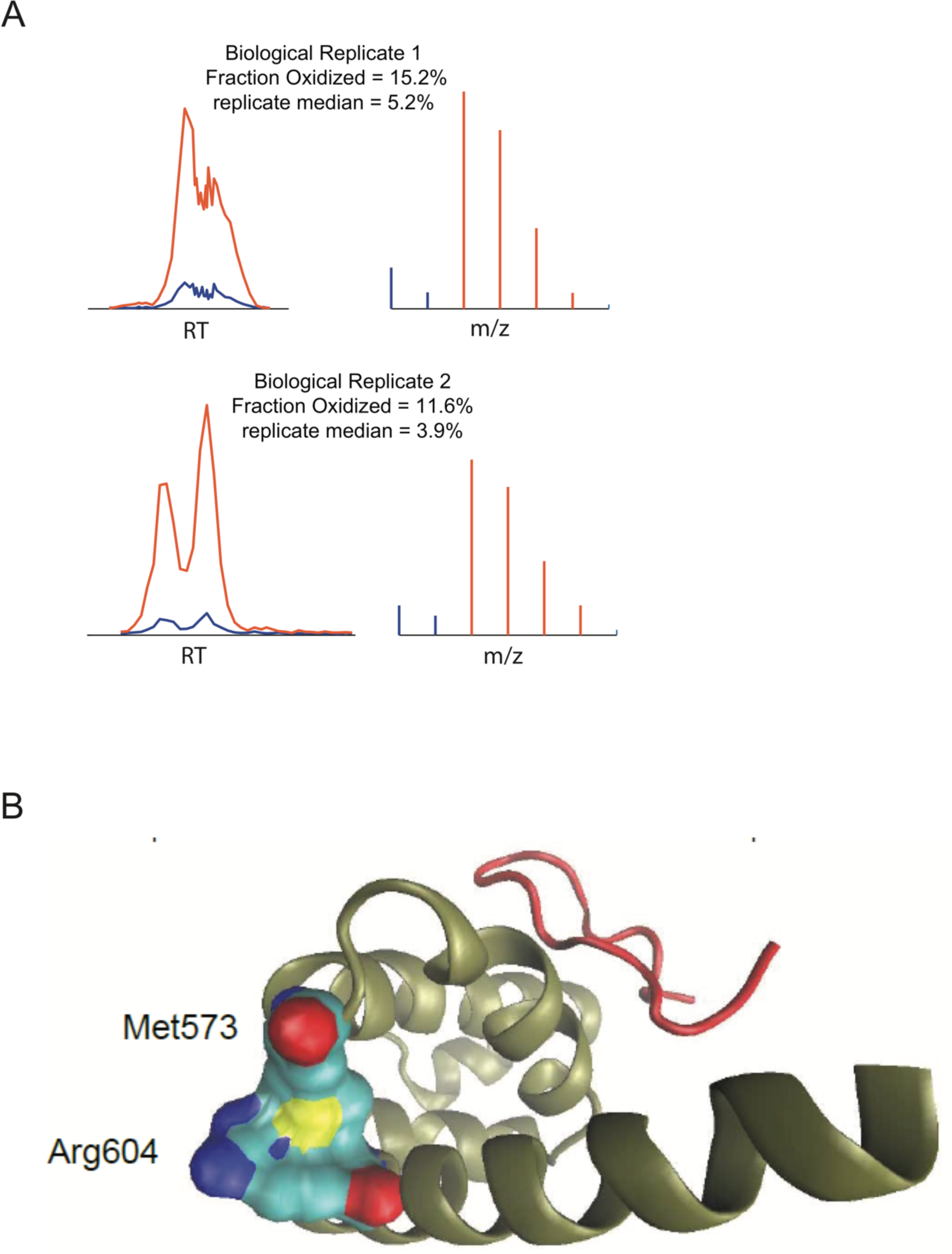
Methionine oxidation of the C-terminal peptide binding domain of PABPC1/3. (A) Models used to calculate the fraction oxidized for the peptide LFPLIQAMHPTLAGK. The total XIC of all features associated with the peptide is shown on the left and the summed MS1 spectra of the best associated feature is shown on the right. Red and blue colors represent the peaks and intensities corresponding to ^18^O and ^16^O peptides, respectively. The “replicate median” values indicate the median fraction oxidized measurements for all methionine-containing peptides in the replicate experiment. (B) The structure of the C-terminal peptide binding domain of PABPC1 (PDB: 3KTP). The interacting partner GW182 is shown in red and the main chain of PABPC1 is shown in gold. Sulfurs are colored yellow, carbons are colored cyan, oxygens are colored red and nitrogens are colored dark blue.

## DISCUSSION

We report a stable isotope labeling strategy that allows for quantification of *in vivo* levels of methionine oxidation on a proteome-wide scale. We used this approach to provide accurate measurements of methionine oxidation in an unstressed human proteome. This methodology relies on labeling and blocking of all *in vivo* unoxidized methionines with a heavy oxygen label immediately following cell lysis. As seen in Figure 2, the labeling and blocking efficiency of methionines was approximately 95%, thereby mitigating the accumulation of *in vitro* methionine oxidation that typically hampers the quantitative analysis of *in vivo* methionine oxidation.

Previous descriptions of the biological significance of methionine oxidation have stemmed from the observation that methionines in proteins are readily oxidized by reactive oxygen species such as hydrogen peroxide. For this reason, methionine oxidation has traditionally been thought of as a form of protein damage that occurs stochastically in response to oxidative stress. This notion, as well as previous measurements of methionine oxidation conducted on oxidatively stressed proteomes, suggested that global levels of methionine oxidation may be moderately high and broadly distributed^17^. However, previous attempts at quantifying global levels of *in vivo* methionine oxidation in unstressed cells have relied on the use of reporter molecules and antibodies with low specificity^21–22^. These approaches may not provide an accurate census of levels of methionine oxidation and do not allow for the quantification of the distribution of methionine oxidation across a diverse array of methionine containing protein sequences. Here, we were able to quantify >3,800 chemically distinct methionine containing peptides in their native context across two biological replicates with ∼1000 peptides shared between them. As seen in Figure 5, baseline levels of *in vivo* methionine oxidation in an unstressed human proteome is tightly distributed around a median value of 5.0% and 3.5% in biological replicates 1 and 2 respectively, which are within the limit of isotopic impurity of ^18^O labeled hydrogen peroxide used as a labeling reagent. In addition, 39.8% and 44.3% of all methionine containing peptides analyzed had levels of oxidation that are below the limit of detection in replicates 1 and 2 respectively. These observations suggest that the levels of methionine oxidation in unstressed cells are maintained at minimal levels and that only a handful of methionine containing peptides are significantly oxidized.

Although the global levels of methionine oxidation were observed to be low, our screen was able to detect a list of peptide sequences having reproducibly high levels of methionine oxidation with high confidence (Figure 5B, Supplementary Table 2). Significantly oxidized peptides were defined using a nonparametric Wilcoxon test comparing the distance from the median for each peptide to the average distance from the median for all peptides. A representative example of a highly confident model that contains direct MS/MS evidence for both the heavy and light peptide and is found to be highly oxidized in both replicates is shown in Figure 6. Met573 from the C-terminal peptide binding domain of polyA binding protein (PABPC1/3) is observed to be approximately 15.2% and 11.2% oxidized *in vivo* in replicates 1 and 2 respectively (p-value=0.023). As can been seen in Figure 6, the sulfur atom of met573 appears to pack tightly against the backbone of Arg604 mediating a spatial interaction between the boundary region of two α-helices. Oxidation of met573 may disrupt this interaction and alter the peptide binding properties of PABPC1/3. Interestingly it has recently been shown that in *Saccharomyces cerevisiae* an interacting partner of PABPC1/3, Ataxin-2, is released from phase separated granules in a methionine oxidation dependent manner in response to mitochondrial stress where it subsequently binds to TORC in order to modulate cellular metabolism and dampen translational activity^23^. Under conditions of mitochondrial stress, the further oxidation of met573 in the peptide binding domain of PABCP1/3 may increase the propensity for Ataxin-2 to interact with TORC by disrupting its interaction with PABPC1/3.

In conclusion, we report a methodology that is able to accurately quantify levels of *in vivo* methionine oxidation across a large subset of methionine containing peptides. The methodology is robustly sensitive toward detecting shifts in the global levels of methionine oxidation and can potentially be used in future studies to monitor changes in the distribution of methionine oxidation levels under various cellular conditions.

## ASSOCIATED CONTENT

The following data are provided in a tabular format in the supplementary information at peptide levels: MS/MS search results and identified peptides, measure fractional oxidation levels and quality control parameters and scores. All raw and processed data are available at ProteomeXchange Consortium via the PRIDE database (accession number PXD014629).

## AUTHOR INFORMATION

### Corresponding Author

*. Phone 585-275-4829

### Author Contributions

The study concept was conceived by J.B. and S.G. Its detailed planning was performed with contribution from all authors. J.B., K.W., and J.H. conducted all experiments. Data analysis was conducted primarily by J.B. with assistance by S.G. The manuscript was written by J.B. and S.G. All authors have given approval to the final version of the manuscript.

### Notes

The authors declare no competing financial interest.

## ACKNOWLEDGMENT

We thank members of the Ghaemmaghami lab at the University of Rochester for helpful discussions and suggestions.

## ABBREVIATIONS

AGC: automatic gain control
BCA: Bicinchoninic Acid
CID: collision induced dissociation
COFRADIC: combined fractional diagonal chromatography
DDA: data dependent acquisition
DTT: Dithiothreitol
DMEM: Dulbecco’s modified Eagle medium
ESI: electrospray ionization
FBS: Fetal Bovine Serum
FDR: false discovery rate
HCD: higher-energy collisional dissociation
IAA: iodoacetamide
MICAL: molecule interacting with CasL
m/z: mass to charge ratio
PSM: peptide-spectrum match
RP-HPLC: reverse phase-high performance liquid chromatography
RT: Retention Time
SDS: sodium dodecyl sulfate
TCEP: tris(2-carboxyethyl)phosphine
TEAB: triethylammonium bicarbonate
TFA: trifluoroacetic acid

## REFERENCES

1. Glaser, C. B.; Li, C. H., Reaction of bovine growth hormone with hydrogen peroxide. Biochemistry 1974, 13 (5), 1044–7.

2. Chao, C. C.; Ma, Y. S.; Stadtman, E. R., Modification of protein surface hydrophobicity and methionine oxidation by oxidative systems. Proc Natl Acad Sci U S A 1997, 94 (7), 2969–74.

3. Samson, A. L.; Knaupp, A. S.; Kass, I.; Kleifeld, O.; Marijanovic, E. M.; Hughes, V. A.; Lupton, C. J.; Buckle, A. M.; Bottomley, S. P.; Medcalf, R. L., Oxidation of an exposed methionine instigates the aggregation of glyceraldehyde-3-phosphate dehydrogenase. J Biol Chem 2014, 289 (39), 26922–36.

4. Hsu, Y. R.; Narhi, L. O.; Spahr, C.; Langley, K. E.; Lu, H. S., In vitro methionine oxidation of Escherichia coli-derived human stem cell factor: effects on the molecular structure, biological activity, and dimerization. Protein Sci 1996, 5 (6), 1165–73.

5. Grintsevich, E. E.; Ge, P.; Sawaya, M. R.; Yesilyurt, H. G.; Terman, J. R.; Zhou, Z. H.; Reisler, E., Catastrophic disassembly of actin filaments via Mical-mediated oxidation. Nat Commun 2017, 8 (1), 2183.

6. Hung, R. J.; Pak, C. W.; Terman, J. R., Direct redox regulation of F-actin assembly and disassembly by Mical. Science 2011, 334 (6063), 1710–3.

7. Lee, B. C.; Peterfi, Z.; Hoffmann, F. W.; Moore, R. E.; Kaya, A.; Avanesov, A.; Tarrago, L.; Zhou, Y.; Weerapana, E.; Fomenko, D. E.; Hoffmann, P. R.; Gladyshev, V. N., MsrB1 and MICALs regulate actin assembly and macrophage function via reversible stereoselective methionine oxidation. Mol Cell 2013, 51 (3), 397–404.

8. Fremont, S.; Hammich, H.; Bai, J.; Wioland, H.; Klinkert, K.; Rocancourt, M.; Kikuti, C.; Stroebel, D.; Romet-Lemonne, G.; Pylypenko, O.; Houdusse, A.; Echard, A., Oxidation of F-actin controls the terminal steps of cytokinesis. Nat Commun 2017, 8, 14528.

9. Balog, E. M.; Lockamy, E. L.; Thomas, D. D.; Ferrington, D. A., Site-specific methionine oxidation initiates calmodulin degradation by the 20S proteasome. Biochemistry 2009, 48 (13), 3005–16.

10. McCarthy, M. R.; Thompson, A. R.; Nitu, F.; Moen, R. J.; Olenek, M. J.; Klein, J. C.; Thomas, D. D., Impact of methionine oxidation on calmodulin structural dynamics. Biochem Biophys Res Commun 2015, 456 (2), 567–72.

11. Kato, M.; Yang, Y. S.; Sutter, B. M.; Wang, Y.; McKnight, S. L.; Tu, B. P., Redox State Controls Phase Separation of the Yeast Ataxin-2 Protein via Reversible Oxidation of Its Methionine-Rich Low-Complexity Domain. Cell 2019, 177 (3), 711–721 e8.

12. Antoine, M.; Boschi-Muller, S.; Branlant, G., Kinetic characterization of the chemical steps involved in the catalytic mechanism of methionine sulfoxide reductase A from Neisseria meningitidis. J Biol Chem 2003, 278 (46), 45352–7.

13. Olry, A.; Boschi-Muller, S.; Branlant, G., Kinetic characterization of the catalytic mechanism of methionine sulfoxide reductase B from Neisseria meningitidis. Biochemistry 2004, 43 (36), 11616–22.

14. Moskovitz, J.; Poston, J. M.; Berlett, B. S.; Nosworthy, N. J.; Szczepanowski, R.; Stadtman, E. R., Identification and characterization of a putative active site for peptide methionine sulfoxide reductase (MsrA) and its substrate stereospecificity. J Biol Chem 2000, 275 (19), 14167–72.

15. Zang, L.; Carlage, T.; Murphy, D.; Frenkel, R.; Bryngelson, P.; Madsen, M.; Lyubarskaya, Y., Residual metals cause variability in methionine oxidation measurements in protein pharmaceuticals using LC-UV/MS peptide mapping. J Chromatogr B Analyt Technol Biomed Life Sci 2012, 895-896, 71–6.

16. Chen, M.; Cook, K. D., Oxidation artifacts in the electrospray mass spectrometry of Abeta Peptide. Anal Chem 2007, 79 (5), 2031–6.

17. Ghesquiere, B.; Jonckheere, V.; Colaert, N.; Van Durme, J.; Timmerman, E.; Goethals, M.; Schymkowitz, J.; Rousseau, F.; Vandekerckhove, J.; Gevaert, K., Redox proteomics of protein-bound methionine oxidation. Mol Cell Proteomics 2011, 10 (5), M110 006866.

18. Rosen, H.; Klebanoff, S. J.; Wang, Y.; Brot, N.; Heinecke, J. W.; Fu, X., Methionine oxidation contributes to bacterial killing by the myeloperoxidase system of neutrophils. Proc Natl Acad Sci U S A 2009, 106 (44), 18686–91.

19. Liu, H.; Ponniah, G.; Neill, A.; Patel, R.; Andrien, B., Accurate determination of protein methionine oxidation by stable isotope labeling and LC-MS analysis. Anal Chem 2013, 85 (24), 11705–9.

20. Shipman, J. T.; Go, E. P.; Desaire, H., Method for Quantifying Oxidized Methionines and Application to HIV-1 Env. J Am Soc Mass Spectrom 2018, 29 (10), 2041–2047.

21. Peterfi, Z.; Tarrago, L.; Gladyshev, V. N., Practical guide for dynamic monitoring of protein oxidation using genetically encoded ratiometric fluorescent biosensors of methionine sulfoxide. Methods 2016, 109, 149–157.

22. Le, D. T.; Liang, X.; Fomenko, D. E.; Raza, A. S.; Chong, C. K.; Carlson, B. A.; Hatfield, D. L.; Gladyshev, V. N., Analysis of methionine/selenomethionine oxidation and methionine sulfoxide reductase function using methionine-rich proteins and antibodies against their oxidized forms. Biochemistry 2008, 47 (25), 6685–94.

23. Yang, Y. S.; Kato, M.; Wu, X.; Litsios, A.; Sutter, B. M.; Wang, Y.; Hsu, C. H.; Wood, N. E.; Lemoff, A.; Mirzaei, H.; Heinemann, M.; Tu, B. P., Yeast Ataxin-2 Forms an Intracellular Condensate Required for the Inhibition of TORC1 Signaling during Respiratory Growth. Cell 2019, 177 (3), 697–710 e17.

